# Comparing 3-D visual and 2-D tactile encounter rates in benthic and pelagic habitats

**DOI:** 10.64898/2026.04.24.720635

**Authors:** Eden Forbes, Jason Stockwell

**Author notes:** Corresponding author: Eden Forbes.

## Abstract

Encounter rate models are important tools for evaluating and estimating trophic interactions between species. While encounter rate parameters have been measured for many freshwater pelagic fishes, most benthic fishes remain mostly unstudied. Those few efforts to generate encounter rate models for benthic fishes often hold mathematical assumptions based on visual foraging, despite the many cases in which benthic fishes employ the lateral line to forage. Furthermore, encounter rate models are rarely compared, despite the many cases in which prey animals face predation risk from multiple types of predators. For example, the macroinvertebrate *Mysis* is exposed to both benthic and pelagic predation risk during diel vertical migration (DVM). Comparing the risks between habitats could help evaluate predation risk as an ultimate cause of their DVM behavior. We created a novel encounter rate model based on lateral line (“tactile”) foraging by sculpins (*Cottidae*) given the saltatory (stop-and-go) nature of their movement. The tactile model demonstrated variation in behavior and peak encounter rate with detection distance, movement velocity, and rest durations. We then directly compared predation risk for *Mysis* by parameterizing both our tactile benthic (2D) encounter rate model for sculpin and a visual pelagic (3D) for rainbow smelt (*Osmerus mordax*). Tactile encounter rates were generally lower than visual rates for individual predators. However, population level encounter rates at night were greater in the benthic habitat than the pelagic habitat. Overall, our model estimates of encounter rates were consistent with the long-standing hypothesis that predation is an ultimate driver of DVM behavior.

## Introduction

Encounters between predators and their prey are critical junctures in the predation cycle and determine trophic interaction strength between species (Lima and Dill 1990, Lima 2002). Models of encounter rates describe both the frequency and probability of encounters given certain physiological and demographic parameters of predators and prey (Koopman 1956, Gerritsen and Strickler 1977). Encounter rate models can be further complexified to account for further biological realism such as patchy prey distributions and variable predator movements (Hutchinson and Waser 2007, Gurarie and Ovaskainen 2013). In lacustrine systems, encounter rate models have been applied to estimate the role of the diel cycle and depth on predation risk (Beauchamp et al. 1999) to describe why foraging strategies shift between prey types and habitats (Persson and Greenberg 1990, Jönsson et al. 2012, Rånaker et al. 2012), and to explain why foraging behavior and/or success changes with conditions such as light, turbidity, and temperature (Table S1).

However, encounter rate models face two major challenges to their broad application in the study of aquatic food webs. First, encounter rate models in lacustrine systems have generally maintained the assumption of ballistic encounters at a fixed and unchanging detection radius, despite the proliferation of encounter rate models that break those assumptions (e.g., Loverdo et al. 2009, Bénichou et al. 2011, Gurarie and Ovaskainen 2013). This is not necessarily an error; for the most part, pelagic freshwater fishes are visual foragers with continuous access to prey information at some distance determined by their physiology and ambient conditions (Guthrie and Muntz 1993). However, assuming visual foraging neglects a large suite of other foraging strategies, such as those that employ the lateral line (Janssen 2004, Mogdans 2019). Benthic fishes especially rely on the lateral line in circumstances where the lack of ambient light precludes efficient visual foraging. Second, encounter rate models are generally only applied with a single foraging species in mind, even though many prey species face risk from more than one type of predator. Navigating multiple kinds of predation risk has significant import on prey distributions and behaviors (Brown et al. 1999; Heithaus et al. 2009; Gaynor et al. 2019). While many studies assess how encounter rates vary within a species of interest, to our knowledge none directly compare encounter rates of two different predator species with the same prey item. Broadening encounter rate models to account for multiple foraging strategies and directly comparing encounter rate models from multiple predators are important steps for employing encounter rates in lacustrine research.

One prescient example in the Laurentian Great Lakes of a prey species that navigates predation risk in benthic and pelagic habitats is the migrating macroinvertebrate *Mysis diluviana* (henceforth, *Mysis*). *Mysis* exhibit diel vertical migration (DVM), moving deeper during the day and migrating up the water column at night (Beeton and Bowers 1982, Gal et al. 2004, Boscarino et al. 2010). The dominant hypothesis is that predator-evasion is the ultimate driver of DVM (Zaret and Suffern 1976, Lampert 1993, Bandara et al. 2019); *Mysis* avoid predation by staying in deeper and darker water during the day and migrate up the water column at night when visual predation risk is low. However, the benthic habitat, historically neglected in discussions of DVM (Stockwell et al. 2020), hosts predators with non-visual foraging strategies such as sculpins (*Cottidae*), which can reliably identify and attack prey in complete darkness (Hoekstra and Janssen 1985, Janssen 1990, Coombs and Conley 1997, Janssen and Corcoran 1998). *Mysis* spend a great amount of time in benthic habitat as part of their DVM behavior and many individuals never leave the benthic substrate (i.e. partial DVM or pDVM; Euclide et al. 2017, O’Malley and Stockwell 2019, Possamai et al. 2025). Consequently, *Mysis* are susceptible to both benthic and pelagic predation over the diel cycle. Directly comparing those risks may help explain *Mysis* distributions and behaviors. While a few DVM models have included both benthic and pelagic predation risk (e.g. Giske et al. 1997, Tarling et al. 2000, Bandara et al. 2018), benthic predation risk is often described as a coarse parameter or with visual foraging assumptions.

We created a novel encounter rate model for a tactile predator in the benthic habitat based on the saltatory (stop-and-go) movement necessitated by sampling the substrate for prey signals (O’Brien et al. 1990, Lewis and Bala 2008, Forbes et al. 2025). We demonstrate how the tactile model behaves according to a variety of factors including prey density, the detection radius or perceptual distance of the predator, and the various movement characteristics that define saltatory motion (movement distance, movement time, and rest time). Then, we compared our newly derived tactile foraging model (2D) to a three-dimensional (3D) visual encounter rate model to compare predation risk for a *Mysis* population from two of their predators: the slimy sculpin *Cottus cognatus* (benthic, tactile predation) and the rainbow smelt *Osmerus mordax* (pelagic, visual predation). To our knowledge, this is the first explicit comparison of encounter rates in 2D and 3D with the same prey species. That said, both the tactile model we derived and our strategy for comparing encounter rate models are generalizable across other habitats and systems. Finally, we call for a greater focus on defining the foraging strategies and encounter rates of benthic fishes.

## Methods

We provide a brief review and amendments to visual encounter rate models that apply to benthic and pelagic environments. Then, we introduce a novel model for tactile foraging in the benthic habitat (2D). In both cases, we are concerned with average destructive encounter rates (Gurarie and Ovaskainen 2013), or the rate of encounters that end in either prey consumption or prey escape from predator detection. We consider both stationary and mobile prey and specifically focus on deterministic encounters, although all the models we describe can be extended to account for probabilistic encounters.

Standard encounter rate models make a consistent set of assumptions based on their derivation from ideal free gas models in physics. First, prey targets are assumed to be dimensionless points in space, although sometimes bestowed with a size characteristic used to determine encounter probabilities (e.g. Beauchamp et al. 1999). Second, prey animals are assumed to be uniformly distributed in space. Third, movement of both the focal predator and the distributed prey species is assumed to be undirected; animals swim in random directions according to a uniform probability distribution. Finally, a predator’s detection radius or reaction distance *d* is often assumed to be uniform in all directions with environmental conditions (e.g. habitat complexity) affecting that distance uniformly. Real detection radii may vary depending on the orientation of the predator’s body (Snyder et al. 2007, Michels et al. 2025), although other studies have found detection radii to be independent of orientation (e.g. Keyler et al. 2021). For the sake of demonstration and comparison between encounter rate models, we will hold all the assumptions above, including uniform detection radii. Consequences of violating some of these assumptions can be found in Hutchinson and Waser (2007) and Gurarie and Ovaskainen (2013).

### Continuous visual foraging model

We first describe model derivations for visual 2- and 3-D encounter rates (Koopman 1956, Gerritsen and Strickler 1977, Hutchinson and Waser 2007). Encounter rates are determined from the predator’s frame of reference. In 2D, encounter rates assume planar polar coordinates with the predator located at the origin and prey targets distributed at various distances and angles from the predator. In 3D, spherical polar coordinates are used, with the predator still located at the origin. In both cases, the predator and individual prey are assumed to be moving at constant velocities in straight lines at random angles with a uniform probability distribution. To be encountered by the predator, a prey animal must enter either the rectangular area (2-D) or cylindrical volume (3-D) that a predator sweeps over while moving. Thus, in 2D, a predicted number of encounters depends on the diameter of the predator’s detection area (2 times detection radius *d*), the mean relative velocity of the prey species to the predator 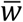, and the observation time *t*. Given some density of prey *P* uniformly distributed in the environment, the number of prey items that fall into the swept area in some interval of time is 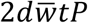 or 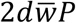 per unit time. Similarly, in 3D the cylindrical volume is defined by mean relative velocity 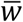 and observation time *t*, this time multiplied by a cross-section of the detection volume (in this case sphere) that intersects the origin (π*d*^2^). The number of encounters with prey density *P* in some interval of time is 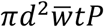 or 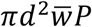 per unit time.

The final ingredient to determine visual encounter rates is an account of mean relative velocity 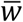. In cases with immobile prey, we can simply replace 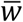 with the mean velocity of the predator 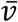, yielding encounter rate formulae with stationary targets for 2D (*E*_*2DS*_; Figure 1A) and 3D(*E*_*3DS*_; Figure 1B):

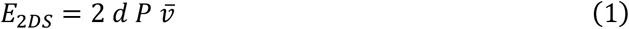

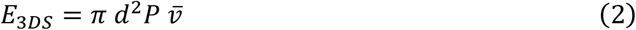

**Figure 1.**
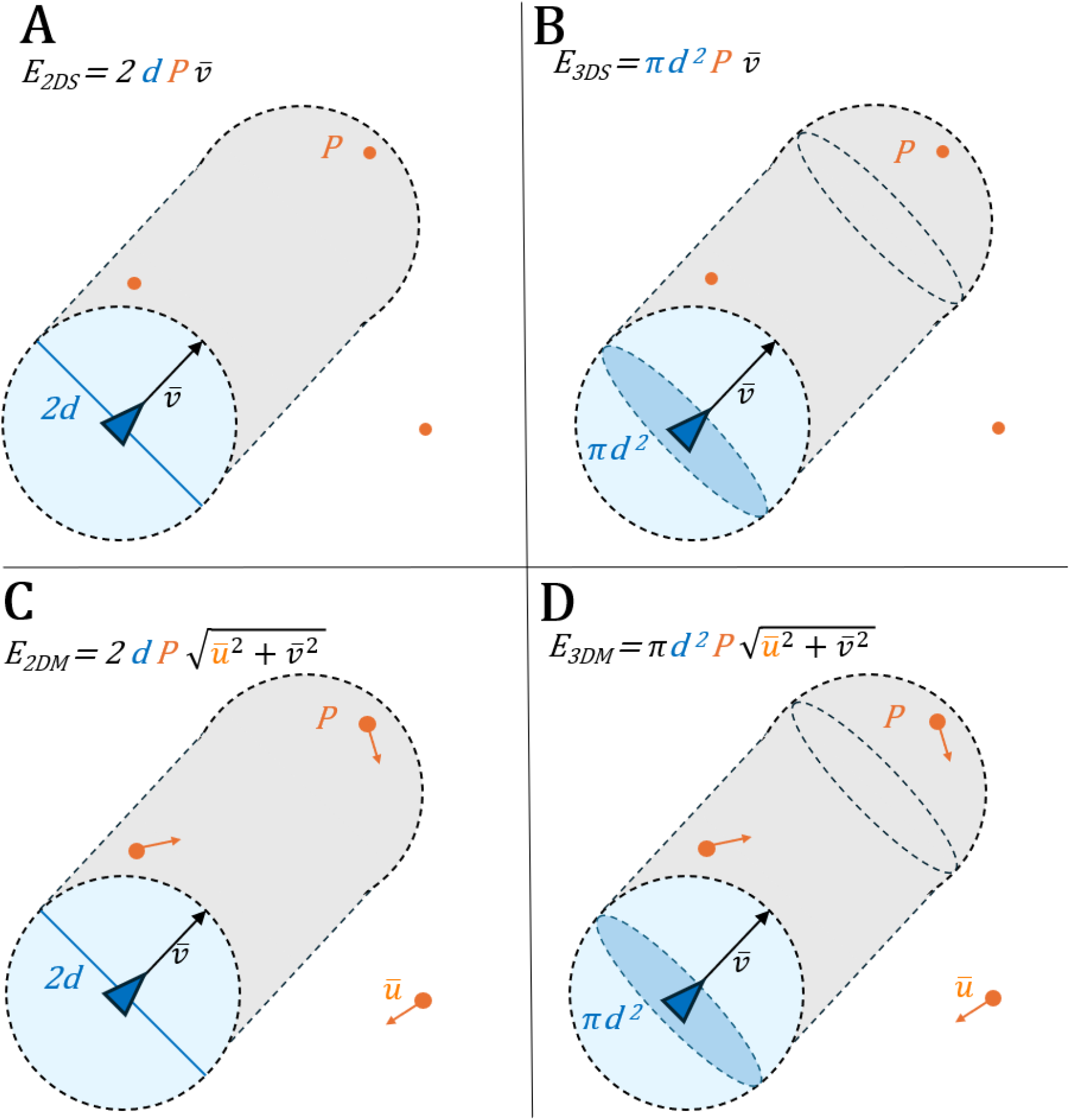
Pictorial representations of the four standard encounter rate models with stationary prey in two dimensions **(A)**, mobile prey in two dimensions **(B)**, stationary prey in three dimensions **(C)**, and mobile prey in three dimensions **(D)**.Blue features pertain to the predator while orange features pertain to prey.

Freshwater ecologists often assume immobile prey to estimate encounter rates (e.g. Beauchamp et al. 1999, Gliwicz et al. 2018) given that zooplanktivore predators move on such a faster scale than their prey that any account of prey movement is unnecessary. That said, modeling encounter rates for piscivores (and at times zooplanktivores, e.g. Pekcan-Hekim et al. 2013) necessitates accounting for prey velocities *u* in the estimation of mean relative velocity 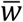.

The velocity of each prey item (*u*) and their predator (*v*) can be combined to determine their relative velocity (*w*) using the law of cosines:

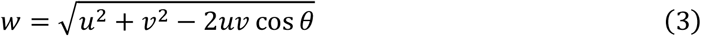

where *θ* is the relative angle between the predator and prey’s trajectories. To determine the mean relative velocity for the entire prey population, the formula can be integrated across a prey velocity distribution. Assuming a uniform distribution of relative predator-prey movement angles (from 0 to π), mean relative velocity 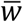 in 2D is given by:

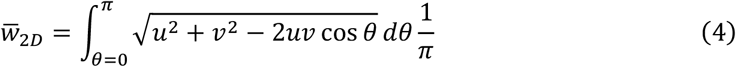

where once again the law of cosines (equation 3) gives the relative velocity for a given relative track angle *θ*. In 3D, the formula is similar, but the distribution of relative predator-prey movement angles is scaled by sin *θ*/2 (Hutchinson and Waser 2007):

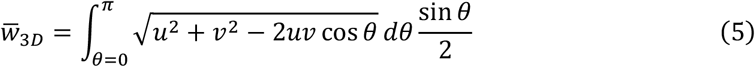

Fortunately, encounter rates can be simplified if the mean movement velocities for predator and prey are known values or follow certain statistical distributions, rather than measured distributions as above. The analytical solution for encounter rates with known mean predator and prey velocities (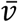 and *ū*, respectively) in 2D and 3D are given by Koopman (1956) and Gerritsen and Strickler (1977), respectively (see also Appendix S2). Each of these solutions can be simplified significantly if predator and prey species have Maxwell-Boltzmann distributions of velocities (Skellam 1958, Evans 1989). In this case, mean relative velocity in both 2D and 3D is given simply by:

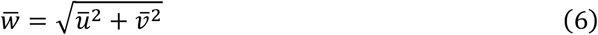

We use this form for mean relative velocity following model comparisons, yielding our final 2- and 3-D encounter rate models (Figure 1C, D):

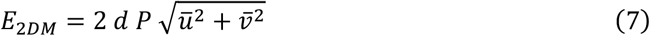

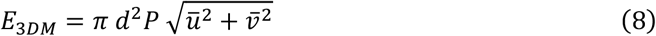

### Tactile benthic foraging model

The encounter rate models described above (equations 1 – 8) are built on a visual model of the detection radius, dating back to their original naval applications (Koopman 1956). That is, equations 1 – 8 assume visual perception at detection radius *d*. If we assumed the visual model applied to benthic predators, encounters would be expected to follow equations 1 and 7. However, many benthic fish rely on tactile stimuli from the lateral line, including signals from motion of the benthic substrate. As a result, some benthic fishes exhibit saltatory motion while foraging on the benthic substrate, with stops to detect prey and movements to search and pursue.

Several excellent models of saltatory behavior with discontinuous sampling exist (e.g., Loverdo et al. 2009; Bénichou et al. 2011; Gurarie and Ovaskainen 2013) but generally rely on numerical simulation (rather than the analytic solutions described for visual encounters above) and sometimes still assume continuous target detection at radius *d*. Here, we introduce a tactile encounter rate model more analogous to *E*_*2DM*_ and *E*_*3DM*_ for the sake of directly comparing encounter rates between benthic (2-D) and pelagic (3-D) habitats with a tactile benthic predator and a visual pelagic predator.

Three variables are necessary to describe the saltatory movement that underlies tactile foraging: the mean displacement of each movement 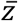, the mean duration of each movement 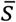, and the mean period of rest between each movement 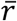. Mean velocity during movement can be given by 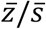 while overall mean velocity can be found by division over the whole saltatory period 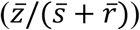. Unlike the previous encounter rate models, the tactile predator does not sweep out a continuous area. Instead, encounters are only possible at rest within period 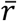. Three cases must be considered to derive tactile encounters with these variables in mind (Figure 2A). First, when the predator is not moving 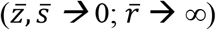 it receives continuous signal given unbroken contact with the substrate. As such, the encounter rate approximates a special case of equation 7 when 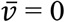:

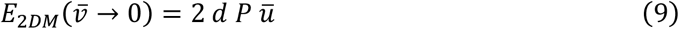

**Figure 2.**
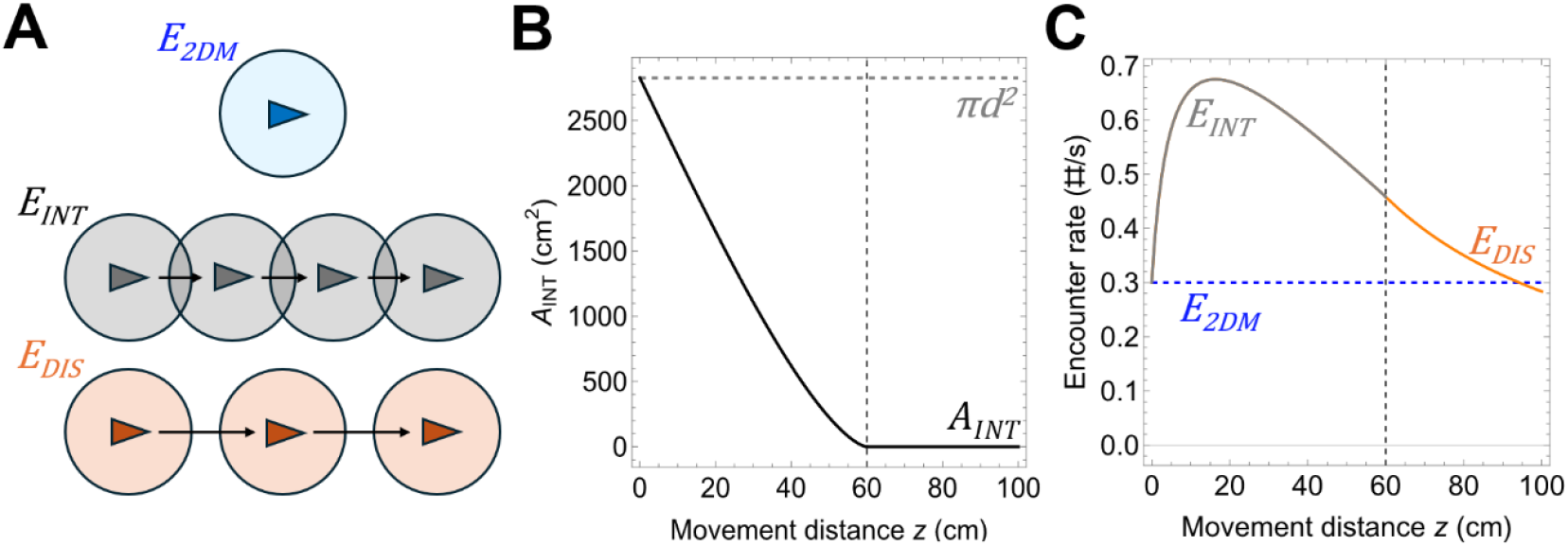
Deriving the encounter rate model for tactile foraging. **(A)** Tactile sampling is not continuous like the visual model except when the predator is at rest (blue). Mobile tactile predators yield two possible situations, either overlapping sensory samples (*E*_*INT*_; gray) or discrete samples (*E*_*DIS*_; orange). **(B)** The amount of overlapping area (*A*_*INT*_; equation 12, black line) in *E*_*INT*_ depends on the mean distance of movement during a saltatory period 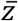. The gray dashed line denotes the total possible sampled area (*πd*^*2*^). **(C)** *E*_*INT*_ generally peaks with some sample overlap but varies significantly depending on model parameters. The y-intercept of *E*_*INT*_ is the same as *E*_*2DM*_, given the functions are identical when the predator is immobile. The dashed black line indicates where the movement distance 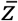 equals twice the detection radius *d*.

Setting 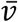 to 0 collapses equation 7 to resemble equation 1, except now the mean velocity of the prey *ū* solely dictates the rate of encounters. Second, consider the opposite extreme in which the predator moves far enough each step that subsequent samples do not overlap 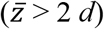. Given that the detection radius *d* is uniform in all directions, the resulting encounter rate with stationary targets (*ū* = 0) is:

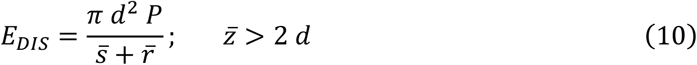

with unique circular samples π*d*^2^ taken every saltatory period 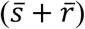. Equation 10 applies if encounters are guaranteed upon contact with the substrate or within rest period 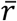. To account for probability of encounter over time while sampling the substrate, encounters would have to be scaled according to some function by mean rest duration 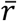. With mobile prey (*ū* > 0), additional encounters can occur during the rest period 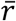. Encounters at rest follow equation 9 multiplied by mean rest duration 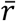, yielding a total encounter rate of:

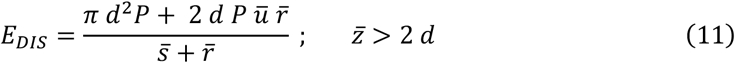

The last case considers predator movement such that sampled areas overlap 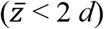. Those areas of intersection must be calculated so previous encounters are not resampled each time the tactile predator comes to rest. The area of intersection (*A*_*INT*_) between two circular samples of radius *d* offset by movement distance *z* can be described as (Figure 2B; Appendix S3):

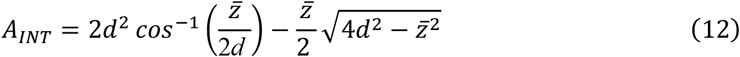

The resampled area described by *A*_*INT*_ is modeled the same as if the tactile predator had never moved – that is according to equation 9 multiplied by the entire saltatory period 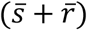. The only encounters counted in the overlapping region are from new prey targets that have entered and settled in that region during the saltatory period. The non-overlapping region is in turn calculated according to the discrete case *E*_*DIS*_ (equation 11). In combination, this yields the intermediate encounter rate (*E*_*INT*_) of:

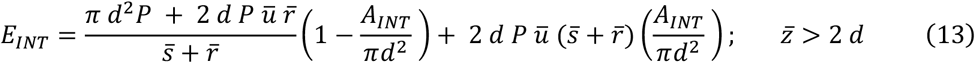

*E*_*INT*_ is defined by equation 13 until the movement distance *z* where the sampled areas do not overlap but also have no space between them 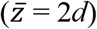, at which point *E*_*INT*_ simplifies to *E*_*DIS*_ (*A*_*INT*_ = 0; Figure 2C). *E*_*INT*_ also simplifies in the case of immobile prey to:

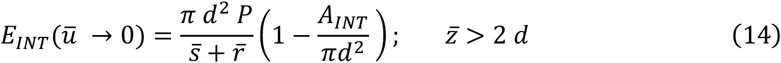

which is the same as equation 10 scaled by the proportion of non-overlapping area sampled.

We assumed that mean movement distance 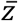 and mean movement duration 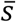 were directly related such that increased 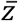 implies increased 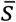. The amount of increase in 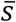 relative to 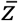 determines the velocity of movement 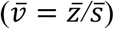. Unless otherwise specified, we assumed that 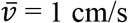.

### Simulations

We assessed the behavior of each encounter rate model (*E*_*2DM*_, *E*_*3DM*_, and *E*_*INT*_) across ranges of predator movement 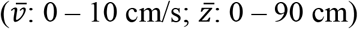. Then, we further characterized the behavior of our tactile foraging model *E*_*INT*_ by varying prey density *P* (0.01 – 0.1 /cm^2^), predator detection radius *d* (1 – 31 cm), mean predator velocity 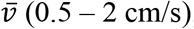, and mean predator rest Periods 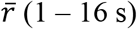. We then compared encounter rates between both benthic (2-D) models (*E*_*2DM*_ and *E*_*INT*_; Appendix S4). Finally, we demonstrated one approach to directly comparing encounter rates between a tactile benthic predator (slimy sculpin; *E*_*INT*_) and a visual pelagic predator (rainbow smelt *Osmerus mordax*; *E*_*3DM*_) given a prey type that is exposed to both kinds of predators (*Mysis*). In this case, we parameterized *E*_*3DM*_ and *E*_*INT*_ according to experimentally verified measures for these species (Table 1) and varied smelt detection radius (*d* = 1 – 18 cm) to proxy for changes from day to night when visual encounter radii are limited. All code used for our model construction and simulation can be found at https://github.com/eforbes24/BP_Encounters.

**Table 1.**
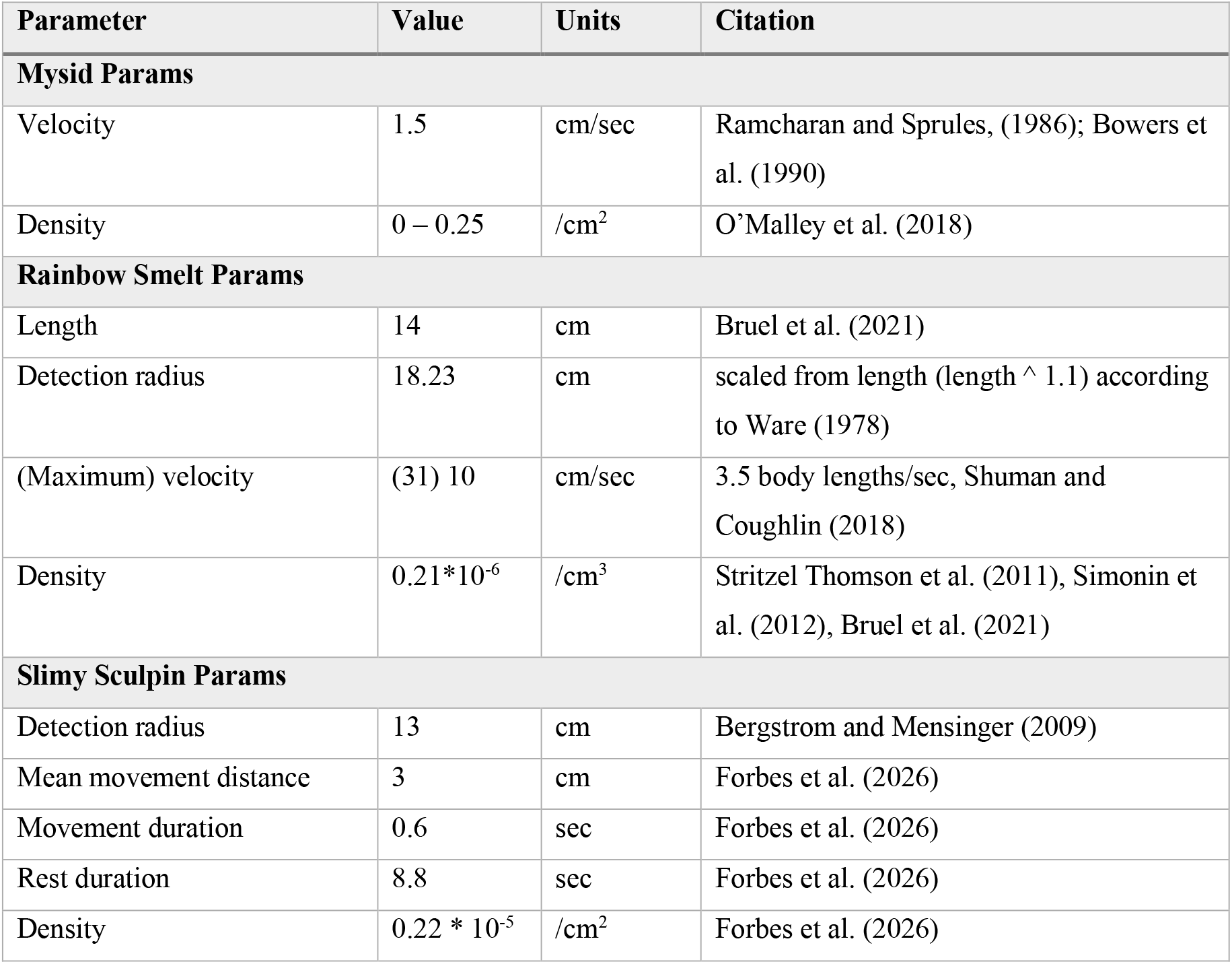
Model notation and parameters used in the derived encounter rate and predation risk models for predators and prey in benthic and pelagic habitats.

## Results

### Visual and tactile encounter rate model behavior

Encounter rates in the standard visual models (*E*_*2DM*_ and *E*_*3DM*_) scaled linearly with predator velocity when prey was stationary (*ū = 0*; Figure 3A, B). A faster moving visual predator encountered more prey, with a doubling of predator swimming velocity resulting in a doubling of the encounter rate. When prey was mobile, encounter rates were primarily determined by the faster of the two species. Encounter rates were more strongly affected by prey velocity at low predator velocities and were entirely determined by prey velocity with stationary predators 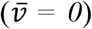. Conversely, as predator velocity exceeded their prey, encounter rates linearly scaled with predator velocity with diminishing impact of prey velocity.

**Figure 3.**
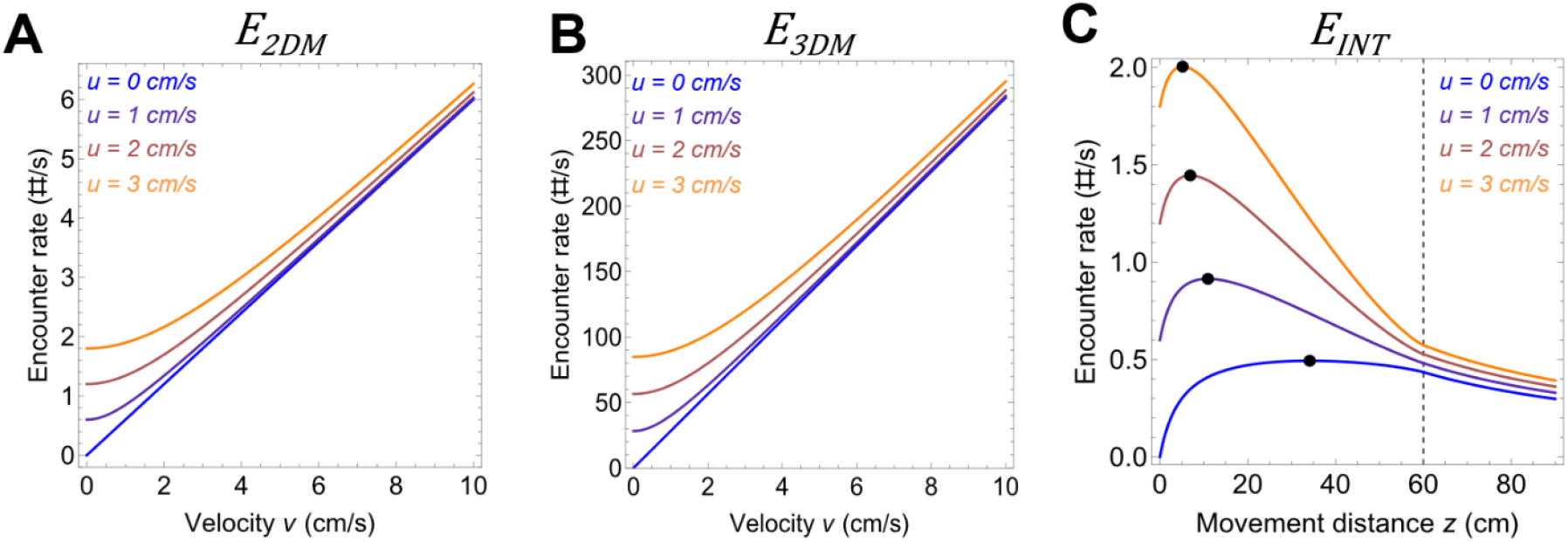
Encounter rates according to predator and prey movement. **(A & B)** Encounter rate for visual predation in 2-D (*E*_*2DM*_) and visual predation in 3-D (*E*_*3DM*_) both as functions of predator velocity 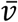 and prey velocity *ū*. Intercepts depend primarily on prey velocity *ū*, but encounter rates converge with greater predator velocity 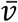 (C) Tactile encounter rates (*E*_*INT*_) in the benthic habitat as a function of predator movement distance 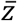 and prey velocity *ū*. The dashed black line indicates the movement distance 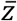 equals twice the detection radius *d*. Black points indicate the predator movement interval 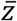 that yields the maximum *E*_*INT*_ at each prey velocity *ū*.

The tactile foraging model *E*_*INT*_ behaved differently than its visual counterparts (Figure 3C). *E*_*INT*_ first increased with increased movement distance 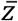 but then decreased as increasing 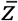 yielded fewer additional encounters per the extra time spent on movement 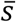. Eventually, increased movement distances no longer yielded any additional encounters as sampled regions no longer overlapped 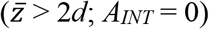. Like *E*_*2DM*_ and *E*_*3DM*_, *E*_*INT*_ was determined by mean prey velocity *ū* when the tactile predator was stationary 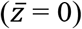. The peak of *E*_*INT*_ was also determined primarily by mean prey velocity *ū*. Increased prey velocity implies a greater proportion of encounters while the predator is at rest as well as greater gains from overlapping samples *A*_*INT*_. Thus, peak *E*_*INT*_ was greater and occurred at lower predator movement distances 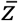 and times 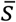 with increased prey velocity *ū*. Differences in *E*_*INT*_ with changing prey velocity became negligible at high predator movement distances and times as time spent in motion 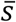 diluted contributions from the rest period 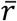.

Like visual encounter rates *E*_*2DM*_ and *E*_*3DM*_, *E*_*INT*_ scaled linearly with increased prey density *P*, yielding greater maximum *E*_*INT*_ but no change in the location of those maxima according to predator movement 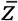 (Figure 4A). Increasing predator detection radius *d* also increased peak *E*_*INT*_, but those peaks shifted to greater predator movement 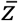 with larger detection radii (Figure 4B). Larger detection radii increased contributions from newly sampled regions from the *d*^*2*^ term from new samples (*E*_*DIS*_, equation 10) relative to those from prey movement while at rest which only scale by *d* (equation 9). However, as parameterized *E*_*INT*_ was far more sensitive to changes in the relative values of predator movement distance 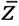, movement duration 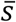, and rest duration 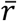 than to detection radius *d*. For example, increasing movement velocity 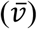 increased the contribution of encounters in newly sampled areas relative to those from prey movement into sampled area (greater increase in [1 – *A*_*INT*_ */d*^*2*^] relative to increases in 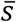). As such, increasing velocity 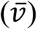 increased both maximum *E*_*INT*_ and pushed those maxima to larger movement distances 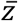 (Figure 4C). Conversely, increasing rest times 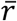 between movements increased the contribution of prey movement into sampled areas. While maximum *E*_*INT*_ decreased with increased rest time 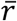, those maxima also occurred at greater movement distances 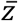 (Figure 4D). Increased rest time increased the contributions of prey velocity to *E*_*INT*_ in both overlapping and newly sampled regions. Thus, by increasing the amount of the saltatory period spent at rest, prey velocity became less important to the optimal movement strategy of the predator, which in turn more closely approximated the optimal movement strategy when prey was immobile.

**Figure 4.**
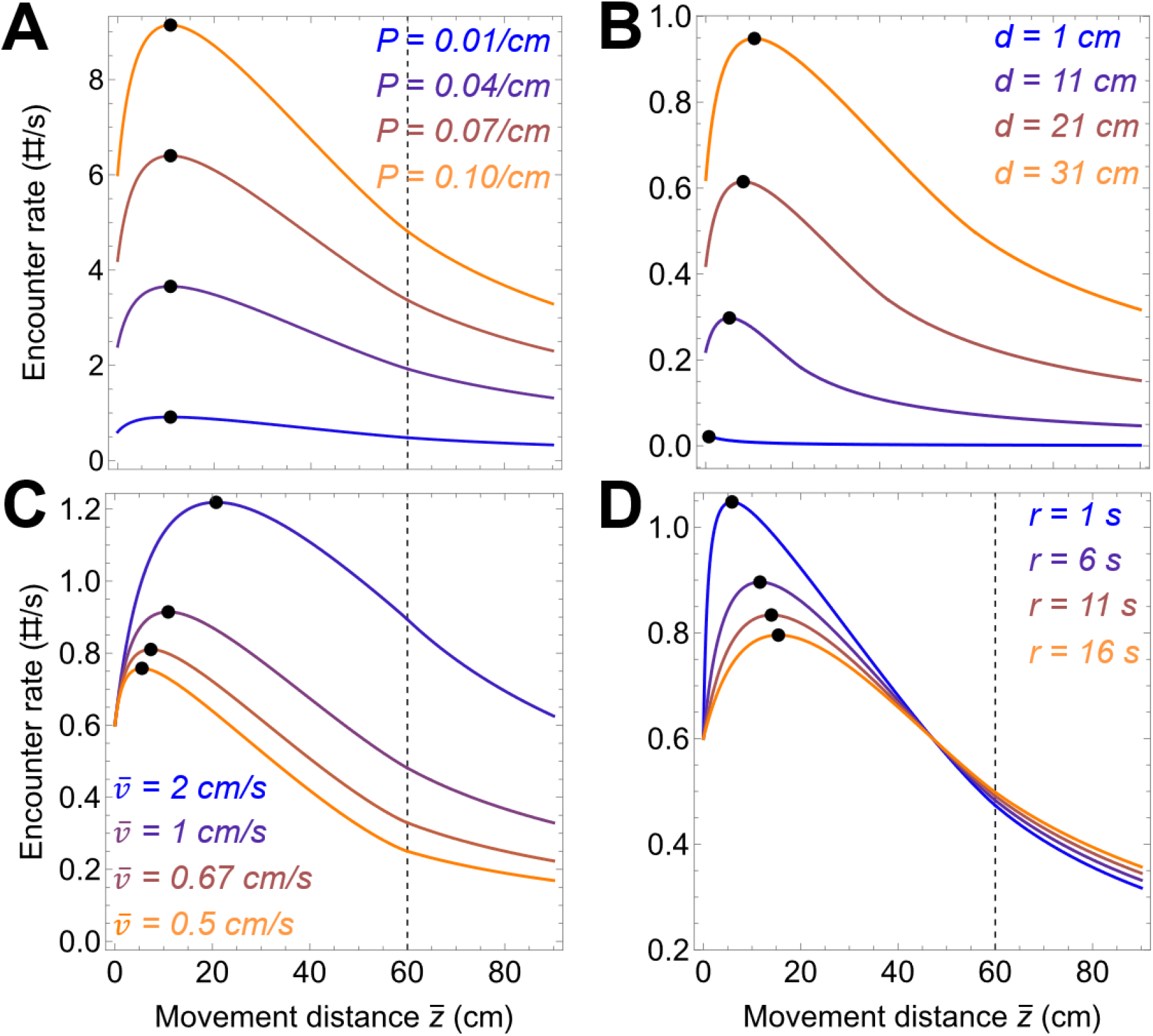
Behavior of the tactile encounter rate model *E*_*INT*_ according to **(A)** prey density *P*, **(B)** detection radius *d*, **(C)** predator velocity during movement 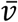, and **(D)** predator rest interval 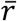. In all cases, the dashed black line indicates the movement distance 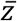 equals twice the detection radius *d*. Black points indicate the predator movement interval 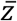 that yields the maximum *E*_*INT*_ according to the relevant parameter in each panel.

### Comparing benthic and pelagic encounter rate models

With equal parameters, the visual models *E*_*2DM*_ and *E*_*3DM*_ yielded greater encounter rates at most predator velocities *v* than the tactile model *E*_*INT*_, even at movement distances 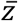 that maximize encounters for *E*_*INT*_ (Figure S2). That said, comparing encounter rates in benthic and pelagic habitats is most useful with specific cases in mind, as the range of possible outcomes is more limited by empirical data. We focused on the example of *Mysis* which negotiates between pelagic predation risk from rainbow smelt and benthic predation risk from slimy sculpin.

Comparing encounter rates in two and three dimensions requires a conversion of areal density to volumetric density; we want a sensible translation of prey density from one habitat to the other. To estimate slimy sculpin and rainbow smelt encounter rates, we had to define a depth of the pelagic habitat those individuals would be distributed within. We made the simplifying assumption that migrating individuals synchronize their DVM and are constrained to a 4-m band of depth that moves up and down the water column. Thus, we examined sculpin and smelt encounter rates in a 1 × 1 m^2^ section of benthic habitat and a 1 × 1 × 4 m^3^ section of the water column, respectively (Figure 5A). Conversion of benthic prey density to pelagic prey density simply required division by 4 (so 1 prey per m^2^ = 0.25 prey per m^3^). Estimated encounter rates for individual rainbow smelt were much greater than that for sculpin during the day, while encounter rates for each species were similar at night when smelt had a reduced detection radius (Figure 5B). At the population level, larger sculpin densities in the benthic habitat yielded population encounter rates that were still lower than that of smelt during the day, but which were greater than that for smelt at night (Figure 5C).

**Figure 5.**
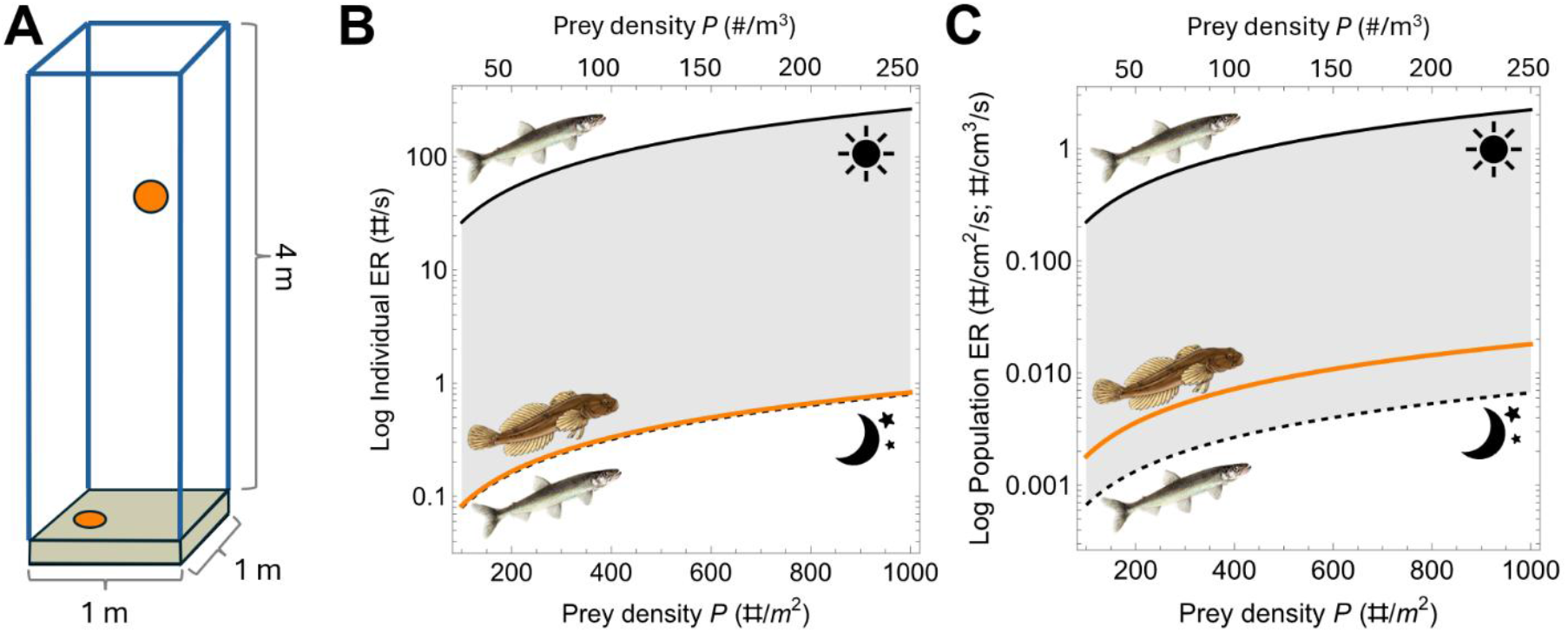
Comparison of encounter rate models in benthic and pelagic habitats. **(A)** Comparing prey densities in benthic and pelagic habitats requires a measure of depth, such that compared densities imply the same number of prey items (orange). Here, we assume 4-m depth in a 1 × 1 m^2^ area. **(B)** Estimated encounter rates for a single slimy sculpin (orange line) and a single rainbow smelt (black lines) according to prey density *P*. Estimated rainbow smelt encounter rates are shown during the day (*d* = 18 cm, solid line) and night (*d* = 1 cm, dashed line). **(C)** Estimated encounter rates for populations of slimy sculpin (orange line) and a single rainbow smelt (black lines) according to prey density *P*. Encounter rate estimates are now given within a unit of area/volume for sculpin and rainbow smelt, respectively. Rainbow smelt encounter rates follow the conventions of (B).

## Discussion

Comparing benthic and pelagic encounter rates requires encounter models that apply to particular species of interest. We have demonstrated a simple approach for comparing encounter rate models between benthic and pelagic habitats by explicitly defining areas and volumes of interest within which to distribute prey populations. We also introduced a new encounter rate model that accounts for the tactile foraging strategy used by many benthic fishes. Our tactile encounter rate model (2-D) used similar movement currencies and parameters as standard (3-D) encounter rate models. Thus, our tactile model is amenable to existing extensions to encounter rate models in the literature including directed movement, non-linear movement, and non-random prey distributions (Hutchinson and Waser 2007, Ioannou et al. 2008, Gurarie and Ovaskainen 2013). Notably, the standard visual encounter rate model sometimes used as a tactile proxy in previous work (e.g., Fiksen and Giske 1995, Giske et al. 1997, Tarling et al. 2000, Bandara et al. 2018) overestimated encounters predicted by the saltatory model even with optimal movement strategies.

Our tactile encounter rate model *E*_*INT*_ varied in behavior from visual encounter rate models in several ways with implications for the study of tactile foragers. First, more mobile prey reduced the movement distances that produced the greatest encounter rates, with rapid prey movement yielding ambush-like strategies. Ambush strategies with high prey velocity are more likely when considering predators also benefit from limiting cost of movement (Huey and Pianka 1981, Killen 2011) and their own ability to be detected by prey. Our models reflect that ambush foraging is one way animals make the best sensory systems that are relatively inefficient or noisy in active search, such as mechanoreception. That said, a variety of animals such as slimy sculpins employ active tactile search and pursuit. Our tactile model suggests that the optimal saltatory movement pattern depends on the velocity of movement used by the predator. High velocities suggest movement intervals that minimize sample overlaps, as predicted by O’Brien et al. (1989), but low velocities suggest significant overlap in samples that maximize *E*_*INT*_. Once again, synthesis of *E*_*INT*_ and energetics models may better reveal what saltatory patterns are efficient for a particular species. Furthermore, saltatory patterns that maximize encounters depend on rest times between samples. While here we assumed that all prey items in the sampled area were detected by the predator, most prey items are behaviorally cryptic, limiting the signals they produce upon their detection of a predator (Ruxton 2009, Stevens and Ruxton 2018). Thus, rest time is a critical variable if longer rest times result in improved detection of cryptic prey. Along those lines, our model suggests that cryptic prey increases saltatory movement intervals that maximize encounters. In all cases we found that the optimal saltatory movement strategy did not change with prey density. Instead, we should expect prey density to primarily affect pursuit and handling time, thus shaping predation pressure but not search strategies.

While encounter rates have been estimated for several individual species, they are rarely compared amongst species, especially with the same prey item in mind. To our knowledge, our application of *E*_*3DM*_ and *E*_*INT*_ to slimy sculpin, rainbow smelt, and *Mysis* is the first direct comparison of 2D and 3D encounter rates in an empirical system. Our application of *E*_*3DM*_ and *E*_*INT*_ recapitulated that *Mysis* benefit from lower pelagic than benthic predation pressure at night, but we also found that benthic predation pressure was greater at night. Our estimate of benthic predation pressure is also conservative, given our trawl data was all from the daytime and sculpin species are caught more frequently at night (Yule et al. 2007). Thus, we expect benthic predation risk to in part drive DVM behavior rather than pelagic resource acquisition alone (Fiksen and Giske 1995, Pearre 2003, Bandara et al. 2018). We suggest that competing predation pressures may shape partial DVM (pDVM) as well. pDVM appears to be ontogenetically mediated, with smaller individuals migrating more frequently than larger ones (Boscarino et al. 2010, Possamai et al. 2025). If smaller *Mysis* benefit from lower detection by pelagic predators (visual) than benthic predators (tactile), then pDVM may be driven by relative predation risk in benthic and pelagic habitats with downstream consequences for benthic-pelagic coupling in lacustrine systems (Chiapella et al. 2023). With direct comparison, encounter rate models can contribute to such larger scale food web questions.

That said, our estimation of predation risk on *Mysis* would be more confident with exact measurement of rainbow smelt detection radius, given we relied on an allometric estimation of smelt detection radius here (Ware 1978). Encounter rate estimates are extremely sensitive to detection radii, thus errors in detection radii estimates are amplified in encounter rate predictions (Vogel and Beauchamp 1999). Fortunately, a bevy of research has targeted variation in detection radii (oftentimes ‘reaction distances’) for Great Lakes fishes (Table S1), including variation in light (Beauchamp et al. 1999, Vogel and Beauchamp 1999, Mazur and Beauchamp 2003, Bergstrom and Mensinger 2009, Keyler et al. 2015, Talanda et al. 2018, Michels et al. 2021, Michels et al. 2025), temperature (Gliwicz et al. 2018), turbidity (Persson and Greenberg 1990, Richmond et al. 2004, Sweka and Hartman 2001, Sweka and Hartman 2003, Rånaker et al. 2012, Peckan-Hekim et al. 2013, Nieman and Gray 2019), and prey type (Howick and O’Brien 1983, Miner and Stein 1996, Jönsson et al. 2012). Encounter rate models have also been extended to explicitly account for some of these environmental factors (e.g. Eggers 1977, Rothschild and Osborn 1988). In combination with swimming velocities during foraging, most of the necessary components to create robust species-specific encounter models are already available for many species. Still, we advocate for more work on the estimation of perceptual ranges and movement behavior of benthic fishes, both of which are far more limited in the literature (although see Hoekstra and Janssen [1985], Janssen [1990], Coombs and Conley [1997], Bergstrom and Mensinger [2009], and Forbes et al. [2026]). Better accounts of benthic foraging are a necessary step to understand the behavior of species in the benthic habitat and those that traverse benthic and pelagic habitats, such as *Mysis*.

More generally, accurate depiction of perception and behavior in any ecological model is critical as behavior is a primary driver of community structure and population dynamics (Martin et al. 2022) and the first response of organisms to changing environments (Wong and Candolin 2015). Constructing appropriate encounter rate models has significant consequences for food web and community-level predictions, such as the cause and magnitude of behaviorally mediated trophic cascades (Werner and Peacor 2003). Behavioral responses of prey are also directly shaped by differential predation risk (Brown et al. 1999; Gaynor et al. 2019), and most often predator-prey populations and distributions are co-determined (Huse and Fiksen 2010; Pinti and Visser 2019) including those that vary on a daily or seasonal basis (Palmer et al. 2017; Kohl et al. 2018) or with habitat complexity (Soukup et al. 2022, Forbes and Stockwell 2026). As such, models of predation should employ perceptually grounded encounter models to unravel the interplay of predator-prey interactions. We suggest that other perceptual modalities, multi-modal encounter strategies, and specific predator-prey relationships be codified in encounter rate models beyond those demonstrated here.

## Acknowledgements

This project was supported by Awards 2018-STO-44074 and 2023-MAR-95003 from the Great Lakes Fishery Commission. Thanks to Randall Beer, Matthew Futia, Olaf Jensen, Amelia McReynolds, Bianca Possamai, and Stephen Thackeray for comments on earlier versions of this manuscript.

## SUPPLEMENTAL MATERIAL

### Section 1: Detection radii for Great Lakes fishes

**Table S1.**
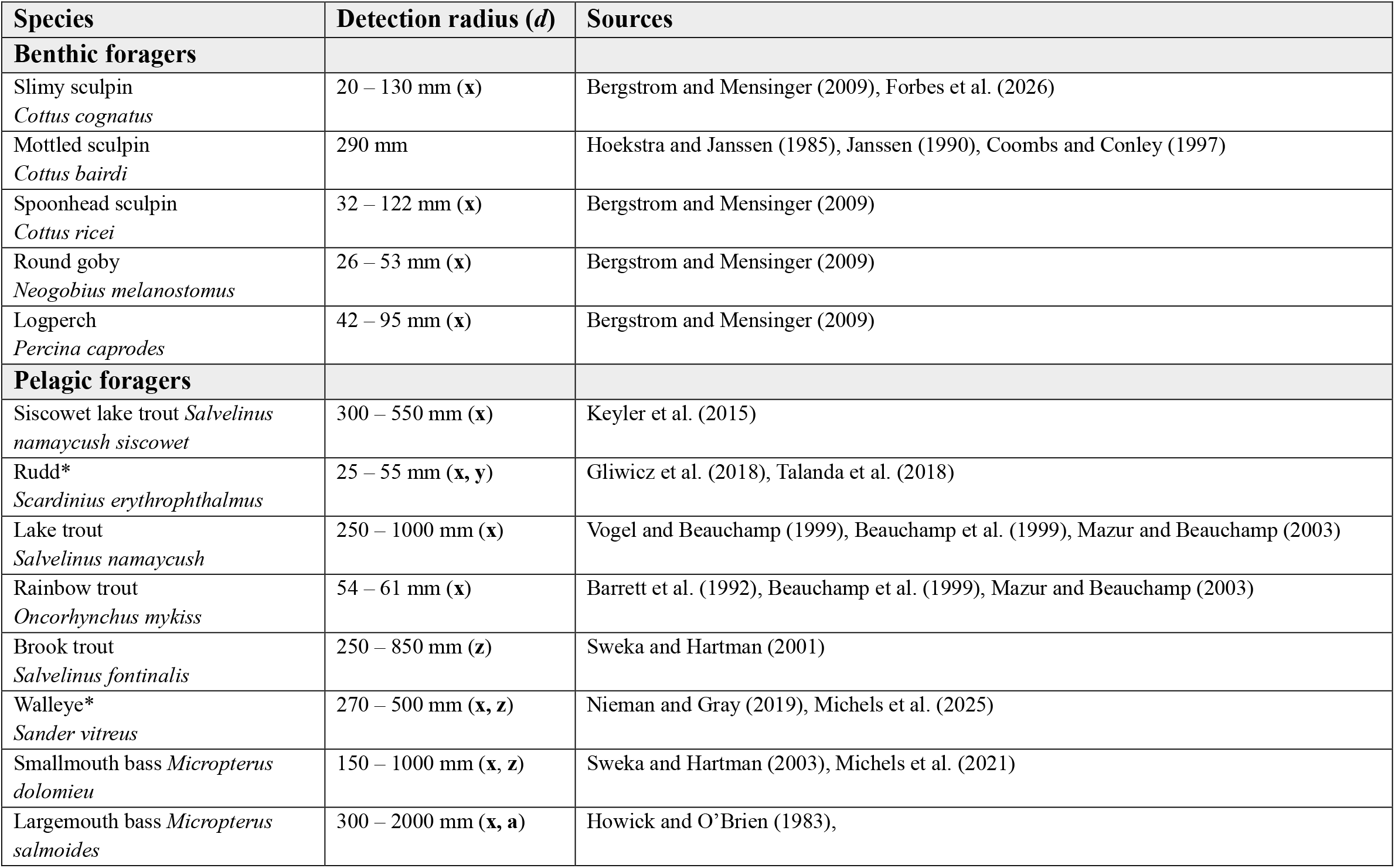
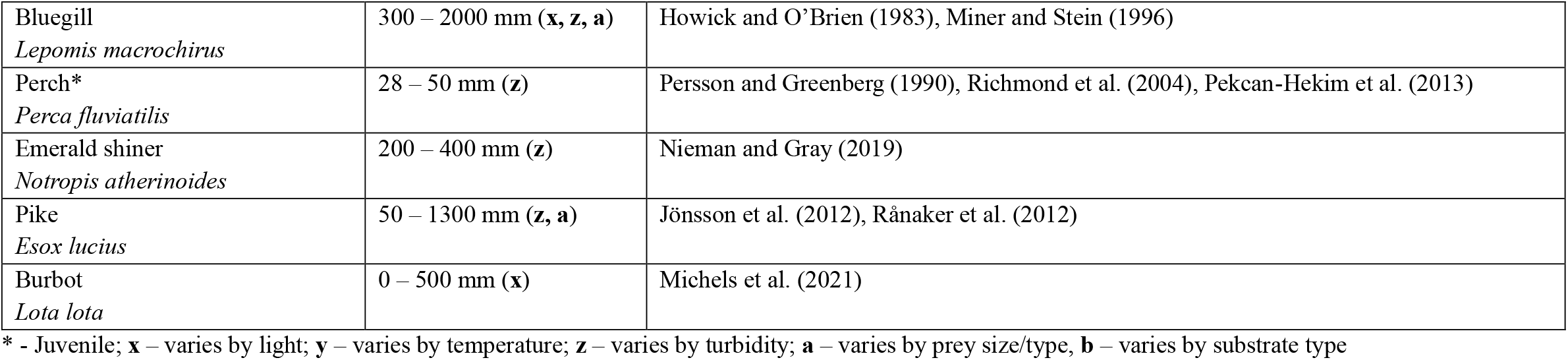
Detection radii from the literature for Laurentian Great Lakes fishes.

### Section 2: Analytical solutions for encounter rates with mobile prey in 2D and 3D

The analytical solution for encounter rates with known mean predator and prey velocities in two dimensions is given by (Koopman 1956):

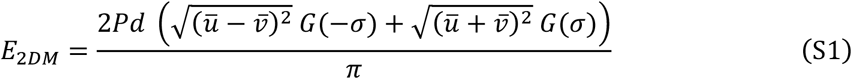

where *G*(*σ*) is Legendre’s complete elliptic integral of the second kind (p. 486 – 487 in Carlson 2010):

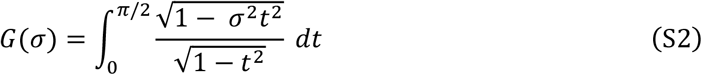

where *t* is the amount of time observed and *σ* determines the number of targets entering the integrated area:

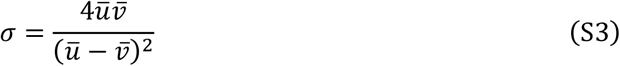

The analogous 3D encounter rate with mobile prey is given by:

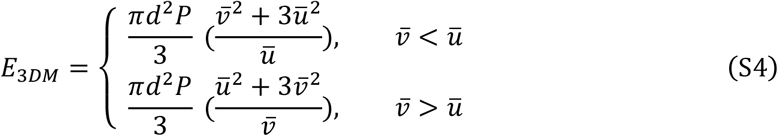

The model is piecewise according to whether the predator or prey species is moving faster but is effectively continuous given it is based on relative velocities between the two species.

### Section 3: Derivation of A_INT_

Determining the area of intersection between two circles is an important part of the saltatory encounter rate model. Fortunately, the problem is simplified given the intersection is symmetric along the axis of movement by the saltatory forager. The distance of each saltatory jump is given by 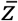. With that distance, we can define the sampled areas as two circles, *C*_*1*_ and *C*_*2*_, with centers offset by 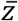 (Fig. S1A):

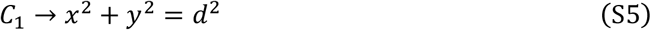

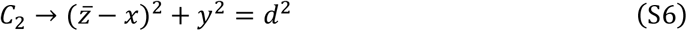

The points (*x, y*) and 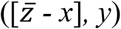 on *C*_*1*_ and *C*_*2*_, respectively, are identical when *x* is half of 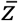. This is one of the points of intersection between the two circles. Knowing the value of *x* is important to determine the area of intersection. To solve for *x*:

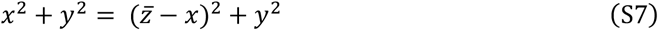

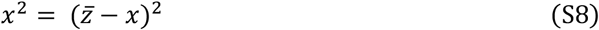

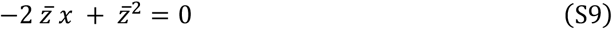

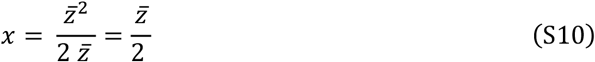

With the value of *x*, we can now solve for the area of the lens through the circle’s center and its points of intersection (connected by chord *a*) with the other circle (shown in Fig. S1B). The area of this lens is given by:

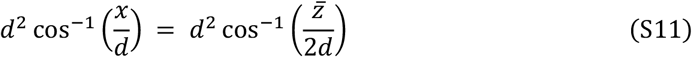

The area of intersection should not account for the triangle below chord *a*, which can be subtracted from this form:

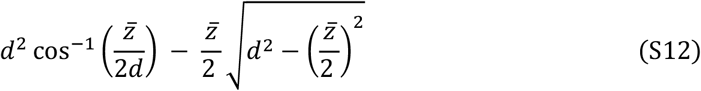

When expanded and doubled (to account for each circle’s lens), we get:

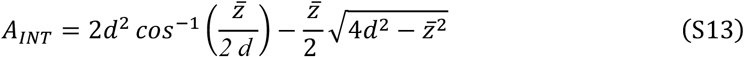

This matches equation 12 in the main text.

**Figure S1:**
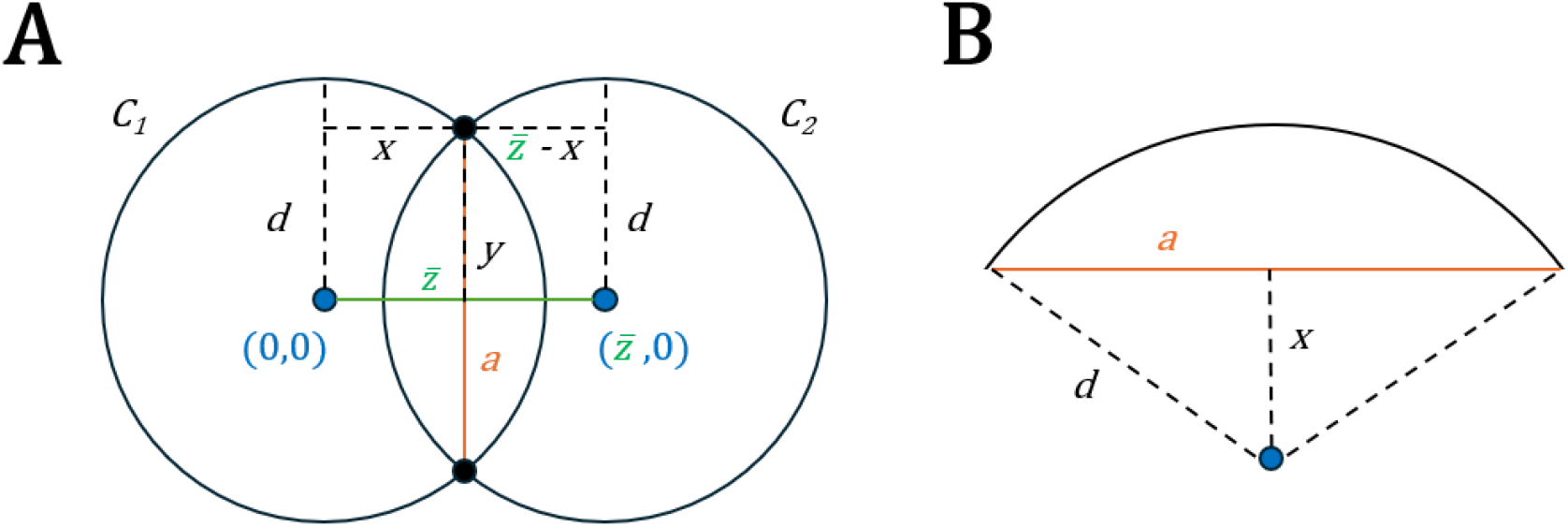
Deriving the area of intersection *A*_*INT*_ between two sampled regions in the saltatory foraging model. **(A)** The area of intersection is defined according to the distance traveled during one saltatory period 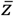. **(B)** Determining the area of the lens defined by that intersection and subtracting the triangle that connects to the circle’s center allows for calculation of the total area of overlap.

### Section 4: Comparison of E_2DM_ and E_INT_

**Figure S2:**
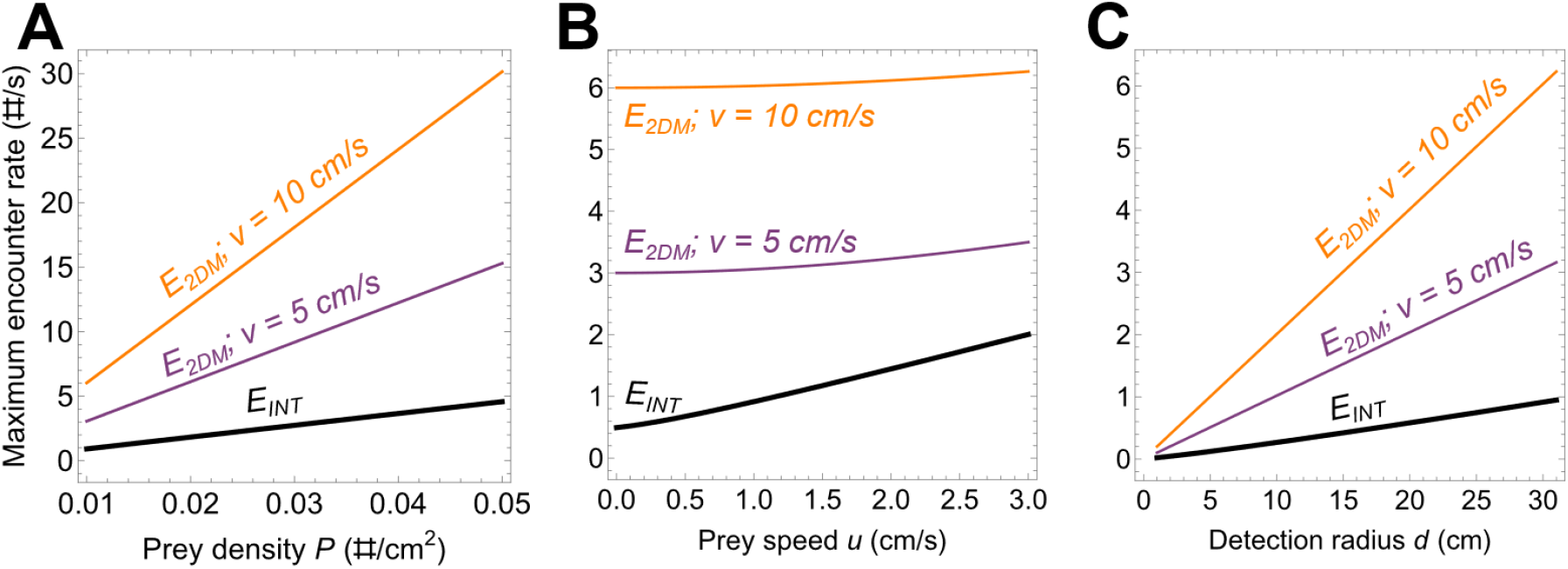
Comparison of *E*_*2DM*_ and *E*_*INT*_ with equivalent parameters varying by **(A)** prey density *P*, **(B)** mean prey velocity *ū*, and **(C)** predator detection radius *d*.

